# A tool to pulse-label yeast Nuclear Pore Complexes in imaging and biochemical experiments

**DOI:** 10.1101/2025.07.15.664992

**Authors:** Annemiek C. Veldsink, Jonas S. Fischer, Sophie Hell, Karsten Weis, Liesbeth M. Veenhoff

## Abstract

Nuclear Pore Complexes (NPCs) are key gateways to the nucleus and major organizers of genome architecture. Despite their importance, it is still not fully understood how NPCs are formed and degraded. Tools to track specific NPCs over time or under stress could unlock critical insights into these questions. Here, we demonstrate that a brief pulse of expression of a previously developed nanobody against baker’s yeast nucleoporin Nup84 (Nordeen et al. 2020) enables a robust, rapid, and straightforward method for pulse-labeling NPCs in both imaging and affinity purification experiments. This approach offers an alternative to permanent yet less rapid genetic fluorophore- or tag-switching techniques and provides a powerful tool for studying NPC inheritance and turnover through both microscopy and biochemical methods.

## Rapid labeling of NPCs using a nanobody against Nup84

Single domain antibodies, nanobodies, have been developed and used for structural biology (Nordeen et al. 2020), for protein purifications (Stankunas and Köhler 2024) and for probing *in vivo* protein assemblies (Solà Colom et al. 2024). A previously developed nanobody (VHH) against the yeast nucleoporin (Nup) Nup84 (Nordeen et al. 2020), a conserved member of the outer ring structures of NPCs, binds Nup84 with nanomolar affinity (VHH-SAN8, from here on VHH[Nup84]) (Nordeen et al. 2020). According to recent structures of the yeast NPC (Singh et al. 2024; Nordeen et al. 2020), the Nup84 epitope is peripherally located in the fully assembled structure, facing both the cytoplasm and nucleoplasm and thus providing an accessible site for nanobody pulse-labeling. To examine the use of VHH[Nup84] as a tool to fluorescently pulse-label NPCs, we fused it to mNeongreen (mNG), a bright fluorophore with short maturation times in yeast (Guerra et al. 2022). We placed the construct under an inducible galactose promoter, allowing for time-controlled expression bursts (Figure 1A), albeit with some cell-to-cell and experimental variability in expression level. Expression of VHH[Nup84]-mNG did not affect cell viability or growth rates, as also previously shown (Nordeen et al. 2020). We observed colocalization of VHH[Nup84]-mNG with NPCs marked by the endogenously tagged NPC component Pom34-mCh (Figure 1B) and no detectable NE signal was observed in nup84Δ cells (Figure 1C), confirming previous structural studies showing that VHH[Nup84] specifically binds Nup84 (Nordeen et al. 2020). Noteworthy, a short, 20-minute expression burst was enough to give a detectable NE signal until at least two hours after the initial induction (Figure 1B, D). Quantifying the signal, we measure that in the first two hours after a 20-minute expression pulse, the mean intensity of VHH[Nup84] reached approximately 46% +/- 27% of the intensity of endogenously tagged Nup84 (Figure 1E). A longer, 1hr-induction pulse further increased the labeling of NPCs compared to a 20-minute expression pulse and the fluorescence levels become indistinguishable (108% +/- 33%) from those obtained with endogenously tagged Nup84 (Figure 1D, E, Figure 1 – supplement 1). Given that the NE labeling intensities are ultimately determined by both incorporation of new VHH[Nup84]-mNG into NPCs as well as the dilution of labeled NPCs to daughter cells, it makes sense that the maximal labeling intensities measured follow the division times in minimal medium (+/- 2hr) (Figure 1E). Collectively, these findings show that pulse-labeling Nup84 in NPCs is quick and tunable, providing an alternative to endogenous tagging of NPC subunits.

**Figure 1.**
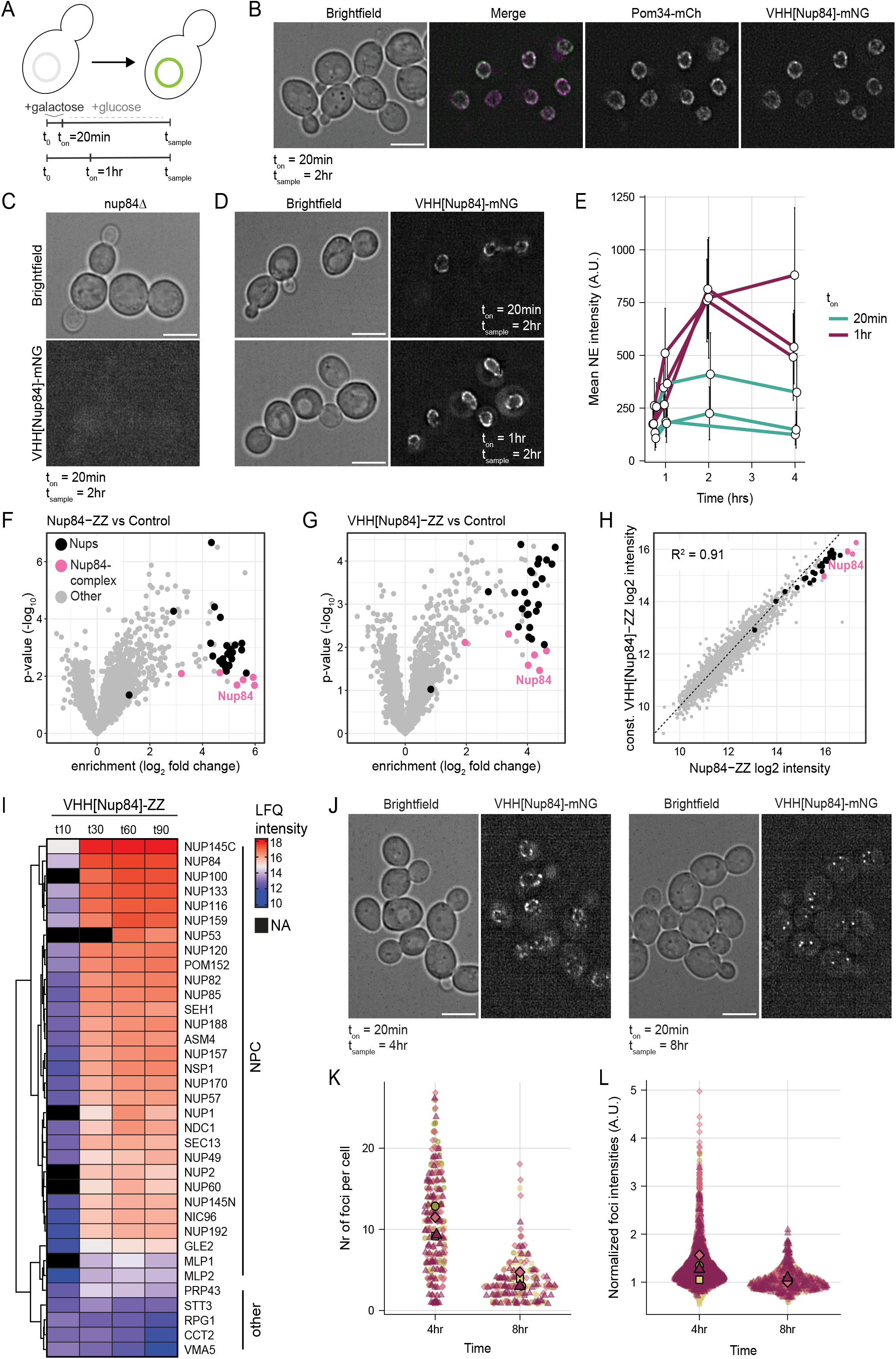
Nanobody against Nup84 is a tool to pulse-label yeast NPCs in imaging and biochemical experiments. **A)** Schematic of the experimental set-up. At t_0_, VHH[Nup84]-mNG is expressed for the specified amount of time (t_on_) by addition of 0.5% galactose to the medium. Expression is subsequently shut off by addition of 1% glucose and images are taken at time (t_sample_) to determine NPC labeling. **B)** Colocalization of VHH[Nup84]-mNG with Pom34-mCh at t=2hr following a 20min induction pulse. Images represent summed projections of 3 z-slices (midplane +/- 0.5 µm). Scale bar = 5 µm. **C)** Typical image of VHH[Nup84]-mNG in nup84Δ cells at t=2hr following a 20min induction pulse. Images represent a single z-slice. Scale bar = 5 µm **D)** Typical images of VHH[Nup84]-mNG at t=2hr following either a 20min or 1hr induction pulse. Images represent sum projections of 3 z-slices (midplane +/- 0.1 µm). Scale bar = 5 µm. **E)** Quantification of NE intensities in the midplane of cells at various timepoints following a 20min (cyan) or 1hr (purple) induction pulse. Data represents mean NE intensity +/- standard deviation of three biological replicas. Numbers of cells analyzed (ton - t_sample_): 102 (20min-45min), 139 (20min-1hr), 82 (20min-2hr, *N=2*), 92 (20min-4hr), 111 (1hr-45min), 186 (1hr-1hr), 153 (1hr-2hr), 193 (1hr-4hr). Mean NE intensity +/- standard deviation of endogenously tagged Nup84-mNG is 726 +/-148 (224 cells). **F)** Volcano plot showing the log2 fold enrichment of proteins in affinity purifications using endogenously expressed Nup84-ZZ as a bait compared to mock-ZZ. Nups are colored in black, the Nup84 subcomplex is colored in magenta. **G)** Volcano plot showing the log2 fold enrichment of proteins in affinity purifications using VHH[Nup84]- ZZ constitutively expressed from the Nup120 promoter as a bait compared to mock-ZZ. Nups are colored in black, the Nup84 subcomplex is colored in magenta. **H)** Correlation between protein intensities in the affinity purifications with VHH[Nup84] from **F** and **G**. **I)** Heat map showing log2 intensities of Nups and five randomly selected non-enriched proteins (“Other”) in VHH[Nup84]-ZZ affinity purifications at the various sampling timepoints following a 20 min VHH[Nup84] induction pulse. LFQ = label free quantitation. **J)** Typical image of the localization of VHH[Nup84]-mNG at t=4hr and t=8hr following a 20-minute induction pulse. Each panel represents a sum projection of 5 consecutive z-slices (0.1 μm). Scale bar = 5 µm. **K)** Quantification of the number of VHH[Nup84]-mNG foci per cell at t=4hr and t=8hr following a 20-minute induction pulse. Each dot represents a cell, each color reflects a biological replica. Means per replica are indicated. Nr of cells analyzed (t_on_ - t_sample_): 289 (20min-4hr), 215 (20min-8hr) from four biological replicas. **L)** Intensities of detected VHH[Nup84]-mNG foci at t=4hr and t=8hr following a 20-minute induction pulse. Intensities are normalized to median foci intensity at t=8hr of the corresponding replica. Each dot represents a single focus, each color reflects a biological replica. Means per replica are indicated. Nr of cells analyzed (t_on_ - t_sample_): 289 (20 min-4 hr), 215 (20min-8hr) from four biological replicas. See Figure 1–Source Data 1 for source data to EFGHIKL.

## VHH[Nup84]-ZZ is a suitable bait for NPC affinity purifications

In addition to the use of VHH[Nup84]-mNG as a pulse-labeling tool in imaging experiments, we characterized the value of VHH[Nup84] as a bait in affinity purifications (APs) of NPCs. We tagged VHH[Nup84] with a tandem Z domain (ZZ-tag), a protein A derived tag that can be purified using IgG-coated beads. We constitutively expressed VHH[Nup84] from the Nup120 promoter and found that all nucleoporins (Nups), except Pom33 and Gle1, significantly coenriched with VHH[Nup84]-ZZ as compared to control APs in cells expressing a free ZZ-tag (**Figure 1G, Source data Figure 1**). Importantly, the level to which Nups are enriched is comparable to APs with endogenously tagged Nup84-ZZ (**Figure 1F**) and overall, there is a strong correlation between APs using the VHH[Nup84]-ZZ and the Nup84-ZZ baits (R2 = 0.91) (**Figure 1H**). This demonstrates that VHH[Nup84] can be used as an affinity bait to purify NPCs, on par with endogenously tagged Nup84-ZZ.

Next, we set out to investigate the affinity purification efficiency of VHH[Nup84]-ZZ when used to pulse-label NPCs. We used a short, 20-minute induction burst to express the VHH[Nup84]-ZZ bait and subsequently determined its interactome at various timepoints (**Figure 1I**). This short expression pulse was enough to specifically purify NPCs 1.5 hours after the induction (**Figure 1I)** and we could already confidently identify most Nups 10 minutes after the start of bait expression. Their intensity increased at the 30 min timepoint and did not drastically change 1.5 hours after labeling start (**Figure 1I)**. This indicates that pulse-labeling is very fast and that VHH[Nup84]-ZZ stays stably bound to the NPC over longer times. Hence, VHH[Nup84]-ZZ can be a powerful tool to pulse-label NPCs to assess the interactome of NPCs at specific points in time.

## VHH[Nup84] can be used to trace labeled NPCs in time

Considering that exchange of individual Nups, including Nup84, is generally slow with half times in the order of hours (Hakhverdyan et al. 2021; Rabut, Doye, and Ellenberg 2004), and that the nanobody:Nup84 association is very stable (Nordeen et al. 2020), we wondered whether we could use VHH[Nup84] pulse-labeling to follow labeled NPCs over longer time periods. Using the same experimental set-up described in Figure 1A, we imaged cells 4hrs and 8hrs after the short 20 min induction pulse. At 8hrs, the signal was diluted from NE rings (Figure 1B, D) to individual foci (Figure 1J), indeed indicating the inheritance of labeled NPCs to daughter cells in mitosis. To quantify the number of foci per cell, we employed the PunctaFinder plugin that can detect foci in 3D (Terpstra et al. 2024). Between 4 and 8hrs, the median number of foci per cell decreased from 11 to 3 (Figure 1K), which is consistent with cell cycle kinetics and a previous reports showing that approximately 40% of the existing NPCs are transmitted to the daughter cell in each division (Zsok et al. 2024; Khmelinskii et al. 2010). The spread in foci intensities decreased in time (Figure 1L), which is likely related to the high density of >10 NPC/µm2 NE (Winey et al. 1997) that compromises the detection of individual pores when they are all labeled. The median intensity reduced by approximately 20% between 4 and 8hrs (Figure 1L), consistent with the stable association of VHH[Nup84] with NPCs. Considering that the rapid labeling of NPCs leads to a quick depletion of unbound VHH[Nup84]-mNG from the cytosol, these foci very likely reflect NPCs that were labeled in the early time stages. Thus, VHH[Nup84]-mNG allows the visualization and tracking of individual NPC over time.

Altogether, we show that VHH[Nup84] is a tool for pulse-labeling of NPCs in live yeast cells in imaging and biochemical experiments. The pulse-labeling of Nup84 in NPCs with VHH[Nup84]-mNG is quick and tunable, providing an alternative to endogenous tagging of NPC subunits. Amongst others this is useful in conditions where direct tagging of Nup84 interferes with its function, while sub stoichiometric nanobody binding may not, or when the temporary introduction of a ZZ- or mNG-tagged nanobody allows assessment of the integrity or localization of mutant NPCs prior to performing experiments without the nanobody. The rapid binding of VHH[Nup84]-mNG to NPCs is stable and hence allows the imaging of individual NPCs in time over multiple divisions. Pulsed expression of VHH[Nup84] thus provides an alternative to the recombination-induced tag exchange (RITE) cassette or optically switchable fluorophores (Verzijlbergen et al. 2010; Zsok et al. 2024; Toyama et al. 2019; Colombi et al. 2013; Zhou and Lin 2013; Onischenko et al. 2020). Lastly, ZZ-tagged VHH[Nup84] can be used as an affinity bait to purify NPCs, on par with endogenously tagged Nup84-ZZ, and, when expressed shortly it can be used to assess the interactome of NPCs at specific points in time or under specific conditions. While our pulse-labeling strategy uses an inducible galactose promoter, other inducible promoters that do not require carbon-source shifts, such as the Gal4-ER-VP16 system (Louvion, Havaux-Copf, and Picard 1993), could be beneficial in conditions when alterations in the metabolic state of the cell are disadvantageous.

As NPCs are long-lived structures, VHH[Nup84] may specifically allow for NPC inheritance studies or the characterization of aged NPCs that were labeled early on during the experiment. Along the same line, we envision that applying genetic perturbations or environmental stresses such as oxidizing conditions, heat stress or osmotic shock at the time of VHH[Nup84] pulse-labeling, provides a means to study the long-term fate of these NPCs. For example, this could resolve the question whether these NPCs are specifically degraded. Similarly, we envision that VHH[Nup84] pulse-labeling strategies can also be used to inducibly bring specific proteins in proximity of NPC, e.g. for determining the local and time resolved interactome of NPCs using proximity labeling strategies (Filali-Mouncef et al. 2024). These and other future studies tracking NPC inheritance and studying NPC turnover mechanisms might uncover how the quality of NPCs is maintained and how this fails in ageing and disease.

## Methods

### Yeast strains and growth

All strains used in this study are listed in the key resources table. Strains were generated using standard microbiology techniques and validated by microscopy and/or PCR. For all pulse-labeling experiments, strains were inoculated 2 days prior to the experiment and grown in synthetic minimal medium supplemented with 2% D-glucose (30°C, 200rpm). On day -1, cells were diluted in minimal medium supplemented with 2% D-raffinose and grown to exponential phase, maintaining them in mid-exponential phase from this point onwards. At the start of the experiment, VHH[Nup84] expression was induced by addition of 0.5% galactose to the medium (2% in the affinity purification experiment; panel I), followed by addition of 1% glucose (2% in the affinity purification experiment; panel I) after 20 minutes or 1hr as described in the figure.

### Fluorescence microscopy

Images were acquired using a Deltavision Elite (Applied Precision) microscope, using an Olympus UPlanSApo 100x (NA 1.4) oil immersion objective and equipped with an EDGE sCMOS5.5 camera for detection. Images were acquired in 30 Z-slices of 0.1 µm, except for panel B (3 z-slices of 0.5 µm). SoftWoRx software (GE Healthcare) was used for image deconvolution and images were processed using Fiji software (ImageJ 2.14.0) (Schindelin et al. 2012). NE intensities were measured in Fiji by drawing line scans over the NE in the midplane and correcting for background intensity. Dumbbell shaped NEs or cells without visible NE staining were excluded from analysis, as were cells in which line scans could not be confidently drawn.

### Affinity purifications and proteomic data acquisition and analysis

Proteomic experiments and data analysis were essentially performed as described in (Kralt et al 2022). To induce expression of VHH bait, cells were grown to mid-log phase as described above and then transferred from medium containing 2% raffinose to medium containing 2% galactose. After 20 minutes of induction, bait expression was halted by directly adding 2% glucose (v/v). At the specified timepoints, samples equivalent to 250 mL of culture of OD600 of 1 were harvested by filtration and snap-frozen in liquid nitrogen. For metabolic labelling experiments, cells were transferred from light L-lysine medium (25 mg/L) containing 2% raffinose to heavy, 13C6 15N2 L-lysine (Cambridge Isotope Laboratories) medium containing 2% galactose to concomitantly start VHH expression and metabolic labelling.

All steps of the affinity purification protocol were performed on ice, minimizing delays to maintain protein complex integrity. Frozen yeast pellets were resuspended in lysis buffer (20 mM HEPES pH 7.5, 50 mM KOAc, 20 mM NaCl, 2 mM MgCl2, 1 mM DTT, 10% v/v glycerol) and transferred to 2 mL screwcap microtubes (Sarstedt Inc.) pre-filled with ∼1 mL of 0.5 mm glass beads (BioSpec Products). The tubes were topped up with additional lysis buffer, and cells were lysed mechanically using a Mini BeadBeater-24 (BioSpec Products) in four 1-minute cycles at 3500 oscillations per minute, with 1-minute cooling intervals in ice water between cycles. Cell debris and unlysed cells were pelleted at 850 x g for 30 s at 4°C and 1 mL of the resulting supernatant was supplemented with 110 µL 10× detergent mix (protease inhibitor cocktail [Sigma-Aldrich], 5% v/v Triton x-100, 1% v/v Tween-20 in lysis buffer). This mixture was transferred to a fresh tube with 1 mg IgG-coated pre-equilibrated magnetic beads (Invitrogen Dynabeads M-270 Epoxy #14301; Sigma rabbit IgG serum #I5006). Samples were incubated at 4°C for 30 minutes with continuous mixing. Beads weore then washed four times with 1 mL of wash buffer (0.1% v/v Tween-20 in lysis buffer) and bound proteins were eluted in 40 µL of 1× Laemmli sample buffer for 2 minutes at 50°C. Eluates were then denatured at 95°C for 5 minutes.

Proteins were concentrated by SDS-PAGE using a 4% acrylamide stacking gel, then incubated in distilled water for 12 hours to remove residual detergents. Protein bands were excised and processed using a standard in-gel digestion protocol. In brief, disulfide bonds were reduced with dithiothreitol (6.5 mM DTT in 100 mM ammonium bicarbonate) for 1 hour at 60°C, followed by alkylation with iodoacetamide (54 mM in 100 mM ammonium bicarbonate) for 30 minutes at 30°C in the dark. Proteins were then digested with 1.25 µg trypsin (Promega) in 100 mM ammonium bicarbonate for 16 h at 37°C. Peptides were desalted using C18 BioPureSPN mini columns (The Nest Group, Inc), washed with Buffer A (0.1% v/v formic acid), eluted with Buffer B (0.1% v/v formic acid, 80% v/v acetonitrile), and ultimately recovered in 12.5 µL Buffer A supplemented with iRT peptides (1:50 v/v, Biognosys).

LC-MS/MS analysis was performed on an Orbitrap Exploris 480 mass spectrometer (Thermo Scientific) coupled to a Vanquish Neo UHPLC system (Thermo Scientific). Peptide separation was achieved on a C18 reversed-phase column (75 μm x 400 mm (New Objective), packed in-house with ReproSil Gold 120 C18, 1.9 μm (Dr. Maisch GmbH)). For the time course samples with the inducible VHH[Nup84] construct, peptides were eluted using a 120-minute linear gradient from 7% to 35% buffer B (0.1% [v/v] formic acid, 80% [v/v] acetonitrile) at a flow rate of 300 nl/min. The mass spectrometer was operated in data-independent acquisition (DIA) mode using one MS1 scan (350-1150 m/z, 120000 resolution, 250% normalized AGC target, 264 ms maximum injection time), followed by 41 variable MS2 windows from 350-1150 m/z with 1 m/z overlap (30000 resolution, 250% normalized AGC target, 64 ms maximum injection time). Fragmentation was carried out by HCD at normalized collision energy (NCD) 28%. For constitutively expressed VHH[Nup84] samples, a 60-minute non-linear gradient from 1% to 43.7% buffer B at a flow rate of 300 nl/min was used. DIA was performed with one MS1 scan (330-1650 m/z, 120000 resolution, 300% normalized AGC target, 20 ms maximum injection time), followed by 30 variable MS2 windows from 330 to 1650 m/z with 1 m/z overlap (30000 resolution, 2000% normalized AGC target, 64 ms maximum injection time). Fragmentation was also performed using HCD, with NCE set to 27%.

The data from the time course samples were extracted with Spectronaut v18.2 (Biognosys) using a spectral library previously generated from 60 APs with 11 Nup baits **(Kralt et al. 2022)**. Data from constitutively expressed VHH[Nup84] were extracted with Spectronaut v19.1 using the directDIA workflow. Carbamidomethylation was used as fixed modification and methionine oxidation, and N-terminal acetylation were set as variable modifications. Tryptic peptides with less than two missed cleavages were considered, and spectra were searched against the *Saccharomyces cerevisiae* protein database (downloaded from SGD on 13/10/2015, 6713 entries). For DIA data analysis in Spectronaut, all default settings were kept except that “cross-run normalization” was turned off. The ion intensities at the fragment level were exported and analysed in R as follows: first, low-quality precursors were excluded based on Spectronaut’s “F. ExcludedFromQuantification” flag. All remaining fragment ions with an intensity > 4 were summed. Only prototypic precursors were retained and precursor intensities were summarized into protein intensities using the MaxLFQ method **(Cox et al. 2014)**. Proteins with less than two identified precursor ions in all three replicates of one timepoint were filtered out. Protein intensities were median normalized per sample, and the mean of three biological replicates was used as the final protein intensity. Statistical significance was assessed using a two-tailed Student’s t-test. The samples with inducible VHH[Nup84] were originally acquired using SILAC with isotopically labelled lysine. However, for our analysis, we relied on label-free quantification by summing the ion intensities of heavy and light y-type fragment ions from precursors containing a single lysine. To account for missing values, pairwise dissimilarities in Fig 1I were computed using the daisy algorithm from the cluster package in R, which accommodates NA values by using all available variable pairs. For comparison, 5 non-NPC proteins were randomly chosen from the list of all reproducibly co-purified proteins. See Figure1-Source Data for LFQ intensities, fold-enrichment, and statistics for all detected proteins.

### PunctaFinder analysis

PunctaFinder analysis (Terpstra et al. 2024) (panels K, L) was essentially performed as described in (Veldsink et al. 2025), using the same threshold settings (Tlocal = 1.478 (ratio punctum/surroundings) and Tglobal = 1.816 (ratio punctum/cell)). Subsequent filtering steps were omitted as the absence of cytosolic VHH[Nup84]-mNG pools did not require additional filtering of detected puncta. Output was analyzed using R software version 4.1.0 (RCoreTeam; 2024). For median normalization of foci intensities (panel L), intensities of all foci at t=4hrs and t=8hrs were divided by the median foci intensity at t=8hr of the same replica.

### Graphs and statistics

Graphs and statistical analysis were performed using R software version 4.1.0 (RCoreTeam; 2024). Data in panels K, L was plotted according to the SuperPlot guidelines (Lord et al. 2020).

## Author contribution

Conceptualization: ACV, JSF, KW, LMV; Investigation: ACV, JSF, SH; Methodology, formal analysis, data curation, visualization: ACV, JSF; Writing—original draft —review & editing: ACV, JSF, KW, LMV; Supervision and funding acquisition: ACV, KW, LMV.

## Acknowledgements

We kindly thank Prof. Dr. Thomas Schwartz (MIT) for sharing the Nup84 nanobody. We thank Federico Uliana and Ino D Karemaker from the IBC MS facility. We thank Michael Chang all members of the Chang and Veenhoff labs for valuable input. Annemiek Veldsink was supported by a PhD-fellowships from the Graduate School of Medical Sciences of the University of Groningen. This work was supported by a Vici grant (VI.C.192.031) to LMV from the Netherlands Organisation for Scientific Research and funding to KW from the Swiss National Science Foundation (TMAG-3_209354 CRSII5_193740).

## Competing interests

The authors declare no competing interests.

## Key Resources Table

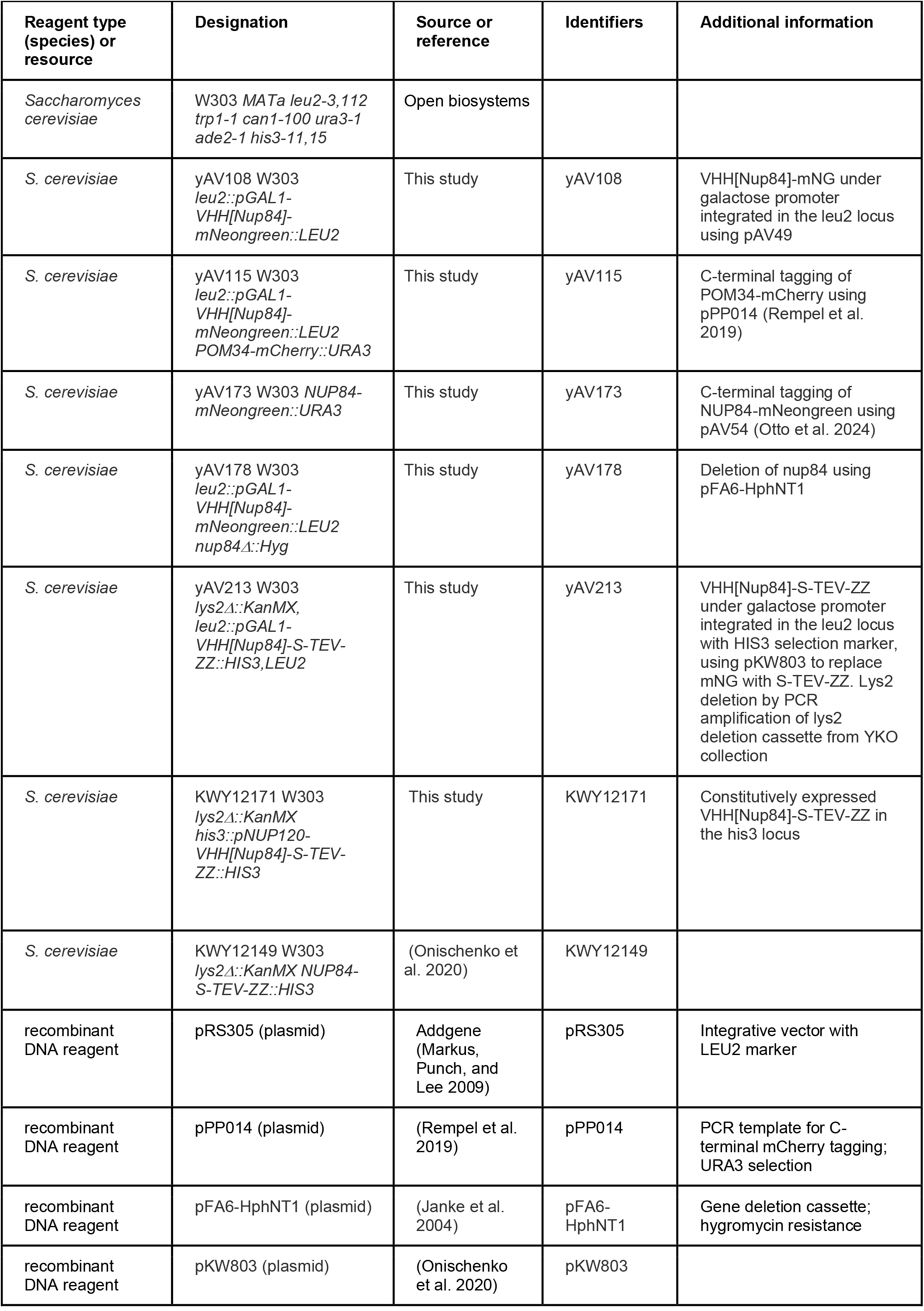
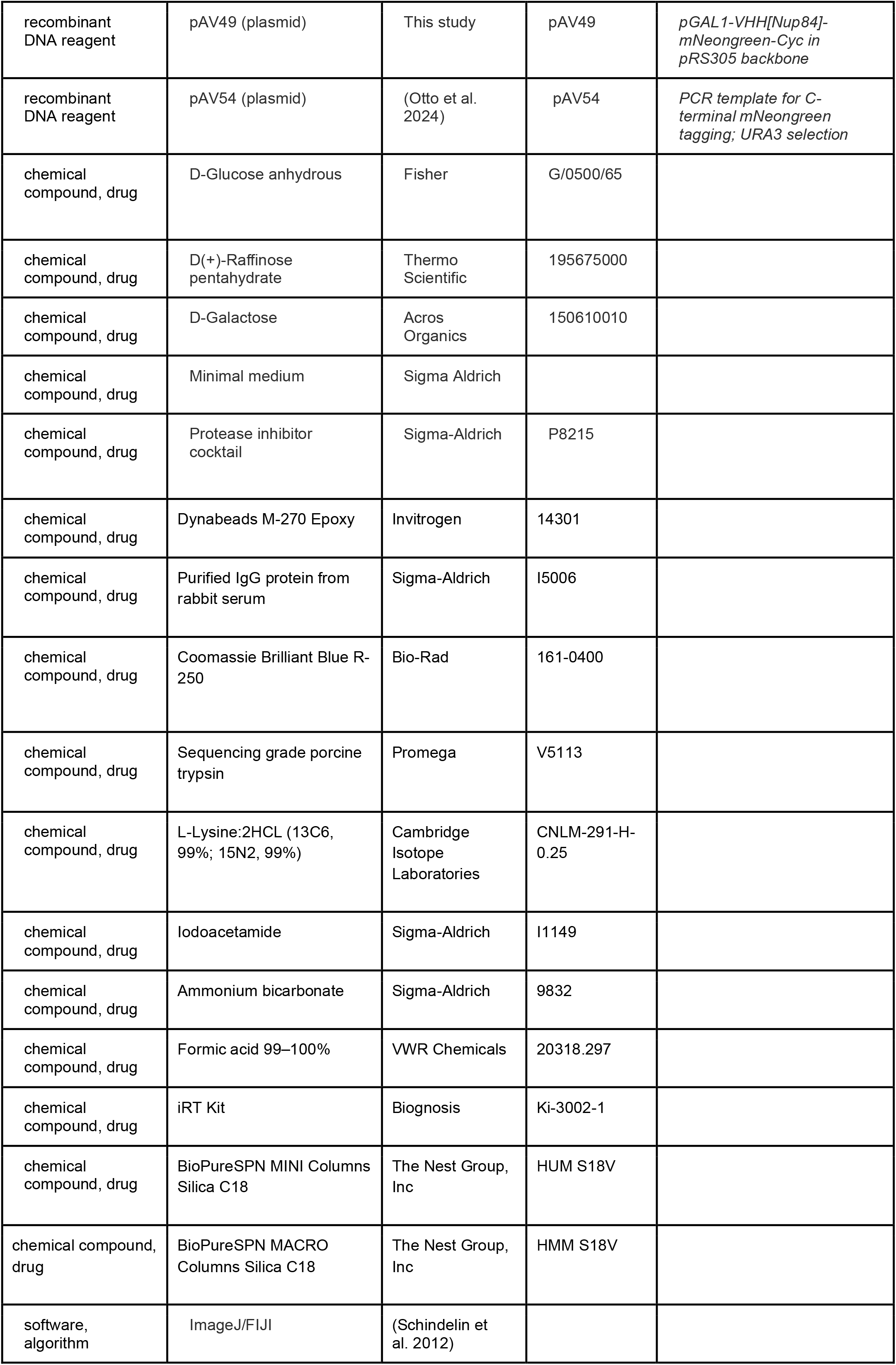
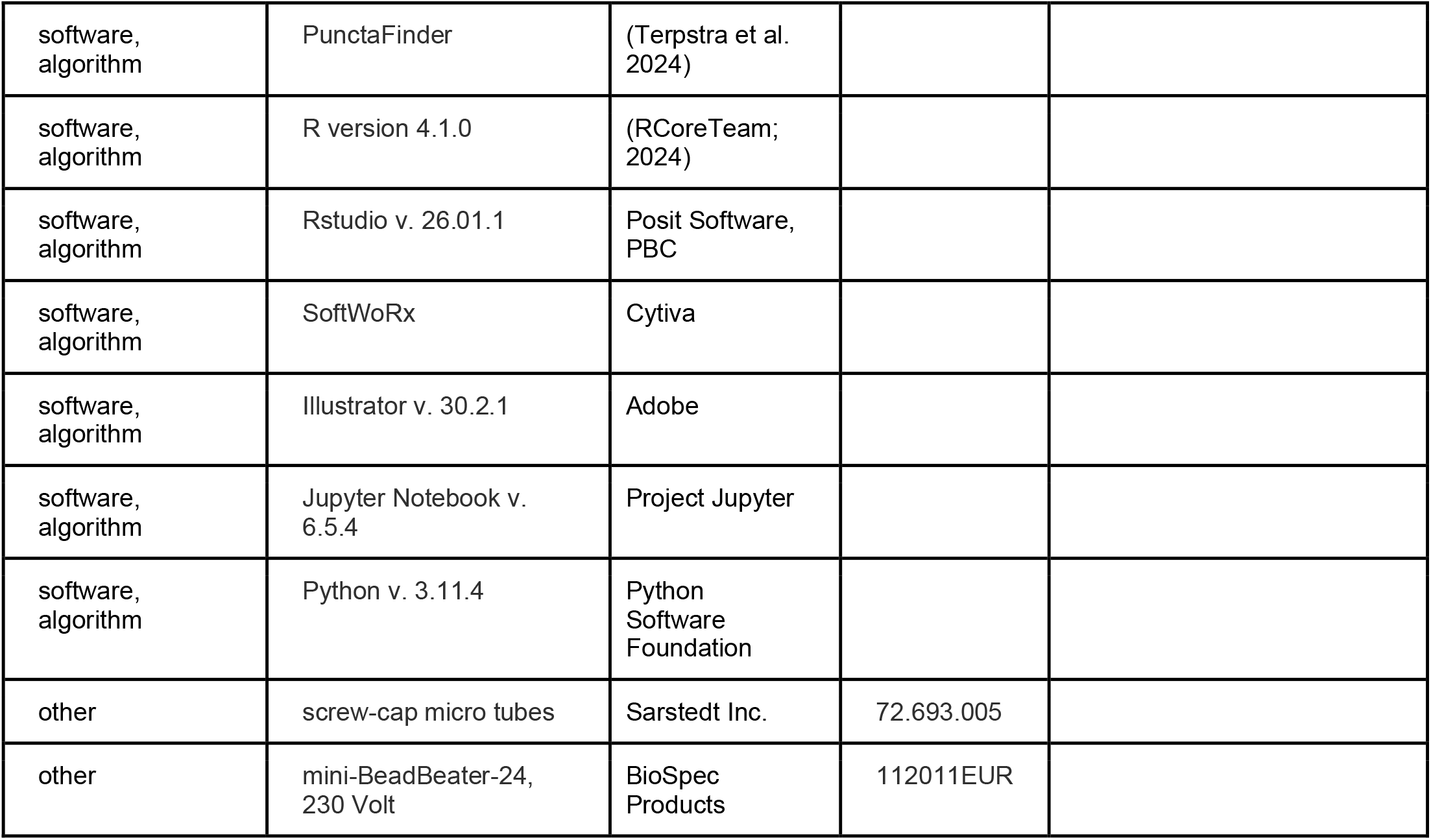

| Key Resources Table |  |  |  |  |
| --- | --- | --- | --- | --- |
| Reagent type (species) or resource | Designation | Source or reference | Identifiers | Additional information |
| *Saccharomyces cerevisiae* | W303 MATa *leu2-3,112 trp1-1 can1-100 ura3-1 ade2-1 his3-11,15* | Open biosystems |  |  |
| *S. cerevisiae* | yAV108 W303 *leu2::pGAL1-VHH[Nup84]-mNeogreen::LEU2* | This study | yAV108 | VHH[Nup84]-mNG under galactose promoter integrated in the *leu2* locus using pAV49 |
| *S. cerevisiae* | yAV115 W303 *leu2::pGAL1-VHH[Nup84]-mNeogreen::LEU2 POM34-mCherry::URA3* | This study | yAV115 | C-terminal tagging of POM34-mCherry using pPP014 (Rempel et al. 2019) |
| *S. cerevisiae* | yAV173 W303 *NUP84-mNeogreen::URA3* | This study | yAV173 | C-terminal tagging of NUP84-mNeogreen using pAV54 (Otto et al. 2024) |
| *S. cerevisiae* | yAV178 W303 *leu2::pGAL1-VHH[Nup84]-mNeogreen::LEU2 nup84Δ::Hyg* | This study | yAV178 | Deletion of *nup84* using pFA6-HphNT1 |
| *S. cerevisiae* | yAV213 W303 *lys2Δ::KanMX, leu2::pGAL1-VHH[Nup84]-S-TEV-ZZ::HIS3, LEU2* | This study | yAV213 | VHH[Nup84]-S-TEV-ZZ under galactose promoter integrated in the *leu2* locus with HIS3 selection marker, using pKW803 to replace mNG with S-TEV-ZZ. *Lys2* deletion by PCR amplification of *lys2* deletion cassette from YKO collection |
| *S. cerevisiae* | KWY12171 W303 *lys2Δ::KanMX his3::pNUP120-VHH[Nup84]-S-TEV-ZZ::HIS3* | This study | KWY12171 | Constitutively expressed VHH[Nup84]-S-TEV-ZZ in the *his3* locus |
| *S. cerevisiae* | KWY12149 W303 *lys2Δ::KanMX NUP84-S-TEV-ZZ::HIS3* | (Onischenko et al. 2020) | KWY12149 |  |
| recombinant DNA reagent | pRS305 (plasmid) | Addgene (Markus, Punch, and Lee 2009) | pRS305 | Integrative vector with LEU2 marker |
| recombinant DNA reagent | pPP014 (plasmid) | (Rempel et al. 2019) | pPP014 | PCR template for C-terminal mCherry tagging; URA3 selection |
| recombinant DNA reagent | pFA6-HphNT1 (plasmid) | (Janke et al. 2004) | pFA6-HphNT1 | Gene deletion cassette; hygromycin resistance |
| recombinant DNA reagent | pKW803 (plasmid) | (Onischenko et al. 2020) | pKW803 |  |
| recombinant DNA reagent | pAV49 (plasmid) | This study | pAV49 | *pGAL1-VHH[Nup84]-mNeogreen-Cyc in pRS305 backbone* |
| recombinant DNA reagent | pAV54 (plasmid) | (Otto et al. 2024) | pAV54 | *PCR template for C-terminal mNeogreen tagging; URA3 selection* |
| chemical compound, drug | D-Glucose anhydrous | Fisher | G/0500/65 |  |
| chemical compound, drug | D(+)-Raffinose pentahydrate | Thermo Scientific | 195675000 |  |
| chemical compound, drug | D-Galactose | Acros Organics | 150610010 |  |
| chemical compound, drug | Minimal medium | Sigma Aldrich |  |  |
| chemical compound, drug | Protease inhibitor cocktail | Sigma-Aldrich | P8215 |  |
| chemical compound, drug | Dynabeads M-270 Epoxy | Invitrogen | 14301 |  |
| chemical compound, drug | Purified IgG protein from rabbit serum | Sigma-Aldrich | I5006 |  |
| chemical compound, drug | Coomassie Brilliant Blue R-250 | Bio-Rad | 161-0400 |  |
| chemical compound, drug | Sequencing grade porcine trypsin | Promega | V5113 |  |
| chemical compound, drug | L-Lysine:2HCL (13C6, 99%; 15N2, 99%) | Cambridge Isotope Laboratories | CNLM-291-H-0.25 |  |
| chemical compound, drug | Iodoacetamide | Sigma-Aldrich | I1149 |  |
| chemical compound, drug | Ammonium bicarbonate | Sigma-Aldrich | 9832 |  |
| chemical compound, drug | Formic acid 99–100% | VWR Chemicals | 20318.297 |  |
| chemical compound, drug | iRT Kit | Biognosis | Ki-3002-1 |  |
| chemical compound, drug | BioPureSPN MINI Columns Silica C18 | The Nest Group, Inc | HUM S18V |  |
| chemical compound, drug | BioPureSPN MACRO Columns Silica C18 | The Nest Group, Inc | HMM S18V |  |
| software, algorithm | ImageJ/FIJI | (Schindelin et al. 2012) |  |  |
| software,<br>algorithm | PunctaFinder | (Terpstra et al.<br>2024) |  |  |
| software,<br>algorithm | R version 4.1.0 | (RCoreTeam;<br>2024) |  |  |
| software,<br>algorithm | Rstudio v. 26.01.1 | Posit Software,<br>PBC |  |  |
| software,<br>algorithm | SoftWoRx | Cytiva |  |  |
| software,<br>algorithm | Illustrator v. 30.2.1 | Adobe |  |  |
| software,<br>algorithm | Jupyter Notebook v.<br>6.5.4 | Project Jupyter |  |  |
| software,<br>algorithm | Python v. 3.11.4 | Python<br>Software<br>Foundation |  |  |
| other | screw-cap micro tubes | Sarstedt Inc. | 72.693.005 |  |
| other | mini-BeadBeater-24,<br>230 Volt | BioSpec<br>Products | 112011EUR |  |

**Figure 1 - supplement 1.**
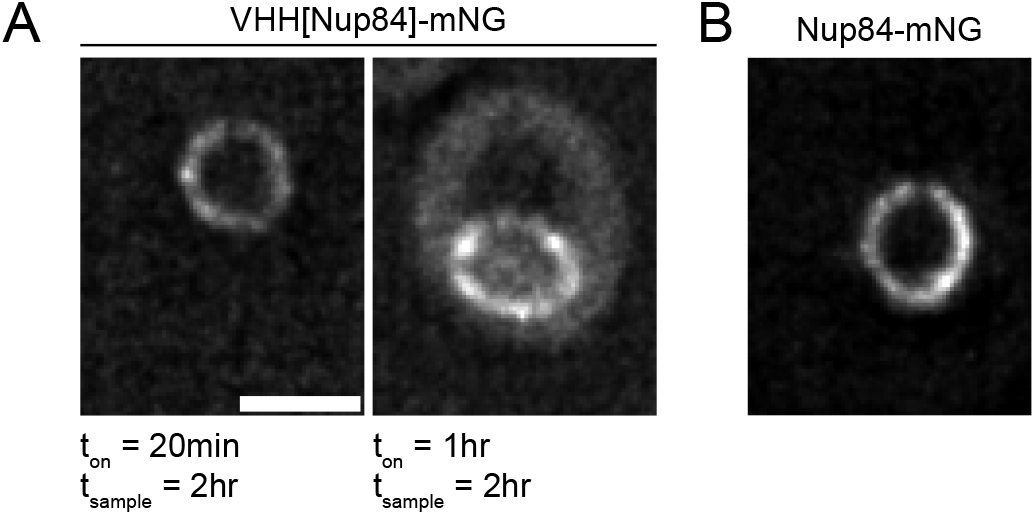
Typical images of a cell expressing VHH[Nup84]-mNG (A) or endogenously tagged Nup84-mNG (B). VHH[Nup84]-mNG expressing cells were imaged at t=2hr following either a 20min (left) or 1hr induction pulse (right). Images represent sum projections of 3 z-slices (midplane +/- 0.1 µm). Scale bar = 2 µm

## Source Data

Figure 1–Source Data 1

## Impact statement

a non-perturbative nanobody-based strategy for pulse-labeling yeast nuclear pore complexes (NPCs) to selectively capture Nup84-containing complexes for imaging and biochemical analysis.

## Notes

### Competing Interest Statement

The authors have declared no competing interest.

### Summary of Updates

Revised version based on comments from eLlife reviewers

